# Review of UNAIDS national estimates of men who have sex with men, gay dating application users, and 90-90-90 data

**DOI:** 10.1101/186163

**Authors:** Reuben Granich, Somya Gupta, Alex Garner, Sean Howell

## Abstract

**Background:** Achieving the 90-90-90 is essential to keep people alive and to end AIDS. Men who have sex with men (MSM) often have the least access to HIV services.

**Purpose:** Estimates for key populations are often unavailable, dated or have very wide confidence intervals and more accurate estimates are required.

**Methods:** We compared registered users from a major gay dating application (2016) from 29 countries with the latest available (2013-2016) UNAIDS estimates by country. We searched the Internet, PubMed, national surveillance reports, UNAIDS country reports, President’s Emergency Plan for AIDS Relief (PEPFAR) 2016 and 2017 operational plans, and conference abstracts for the latest nationally representative continua for MSM.

**Results:** Of comparison countries, only 18 countries had UNAIDS or other MSM population estimates in the public domain. UNAIDS estimates were larger than the gay dating application users in 9 countries, perhaps reflecting incomplete market penetration for the application. The gay dating application users in 9 countries were above the UNAIDS estimates; 8 were over 30% higher and three more than double the reported estimate. Seven partial or complete nationally representative care continua for MSM were published between 2010 and 2016. Among estimated MSM living with HIV, viral suppression varied between 42% (United States) to 99% (Denmark). The quality of the continua methods varied (quality data not shown).

**Conclusion:** “What is not monitored is not done” and social media has significant promise to improve estimates to ensure that MSM and other vulnerable people living with HIV and their communities are not left behind on the way to ending AIDS.

## Introduction

More than 37 million people are living with HIV with 18.2 million (50%) on treatment that prevents illness, death and transmission (1). Achieving the 90-90-90 target (By 2020, 90% of all people living with HIV will know their HIV status, 90% of all people with diagnosed HIV infection will receive sustained antiretroviral therapy, and 90% of all people receiving antiretroviral therapy will have viral suppression) is essential to keep people alive and to end AIDS (2). Men who have sex with men (MSM^1^) bear disproportionate burdens of HIV but often have the least access to HIV services. Stigma and discrimination remain as major factors in MSM vulnerability to HIV and lack of access to services. Although population estimates are available for adults and children, estimates for key populations including men who have sex with men (MSM) are often unavailable, dated or have very wide confidence intervals (3). Estimates of MSM are rarely population-based or representative and often rely instead on potentially biased smaller observational studies or surveys in sexually transmitted disease clinics or other service settings (4-9). The lack of accurate MSM population estimates compromises the HIV response in a number of important ways including the ability to plan and budget for MSM specific services, ability to measure progress in service delivery and health impact, and impaired ability to address stigma and discrimination. The dearth of solid denominators also makes it problematic to estimate the 90-90-90 target that is vital for preventing illness, death and transmission among MSM.

Over 3.5 billion people worldwide are connected to the Internet with over 2 billion from developing countries‐‐cloud-based information and applications are increasing important in everyday life (10). Public health has warily embraced social media and there are an increasingly wide array of applications to solve public health challenges (11-14). Calls to improve individual and public health through open data, crowd-sourcing information and big data have been met with general enthusiasm although some have urged caution. Social media increasingly reflects and influences and reflects the sexual habits of millions of people who use applications for meeting people, dating, and to arrange to have sex. With millions of users coming from a wide spectrum of the society, social media users may provide new insights into population size estimates.

We used data from a popular gay software application that is designed to make it easy for gay, bi, and curious guys to connect in various ways (15). It is a global application and has over 20 million unique users in over 30 countries. Its widespread popularity, and relative anonymity raises the question whether the number of users can be used to improve official the Joint United Nations Programme on HIV/AIDS (UNAIDS) and national MSM estimates.

## Methods

To examine the relationship between the gay dating application users and MSM population estimates, we compared the 2016 registered dating application users from 29 countries with the latest available (2013-2016) UNAIDS (16) and national estimates (6, 17) by country (Figure 1). Countries were selected with over 27,000 gay dating application users to protect their identity and possible identification. Only unique individual dating application users active during the study period were identified. We applied an algorithm that uses device, location, usage, and registration information to ensure that only unique individual dating application users were counted. Methods to determine the UNAIDS estimates for MSM rely on observational studies, surveys and expert opinion. To address progress towards the 90-90-90 target and as an insight to access to services, we also searched the Internet, PubMed, national surveillance reports, UNAIDS country reports, President’s Emergency Plan for AIDS Relief (PEPFAR) country/regional operational plans 2016 and 2017, and conference abstracts for the most recent, complete and nationally representative continua for MSM (18-20).

**Figure 1.**
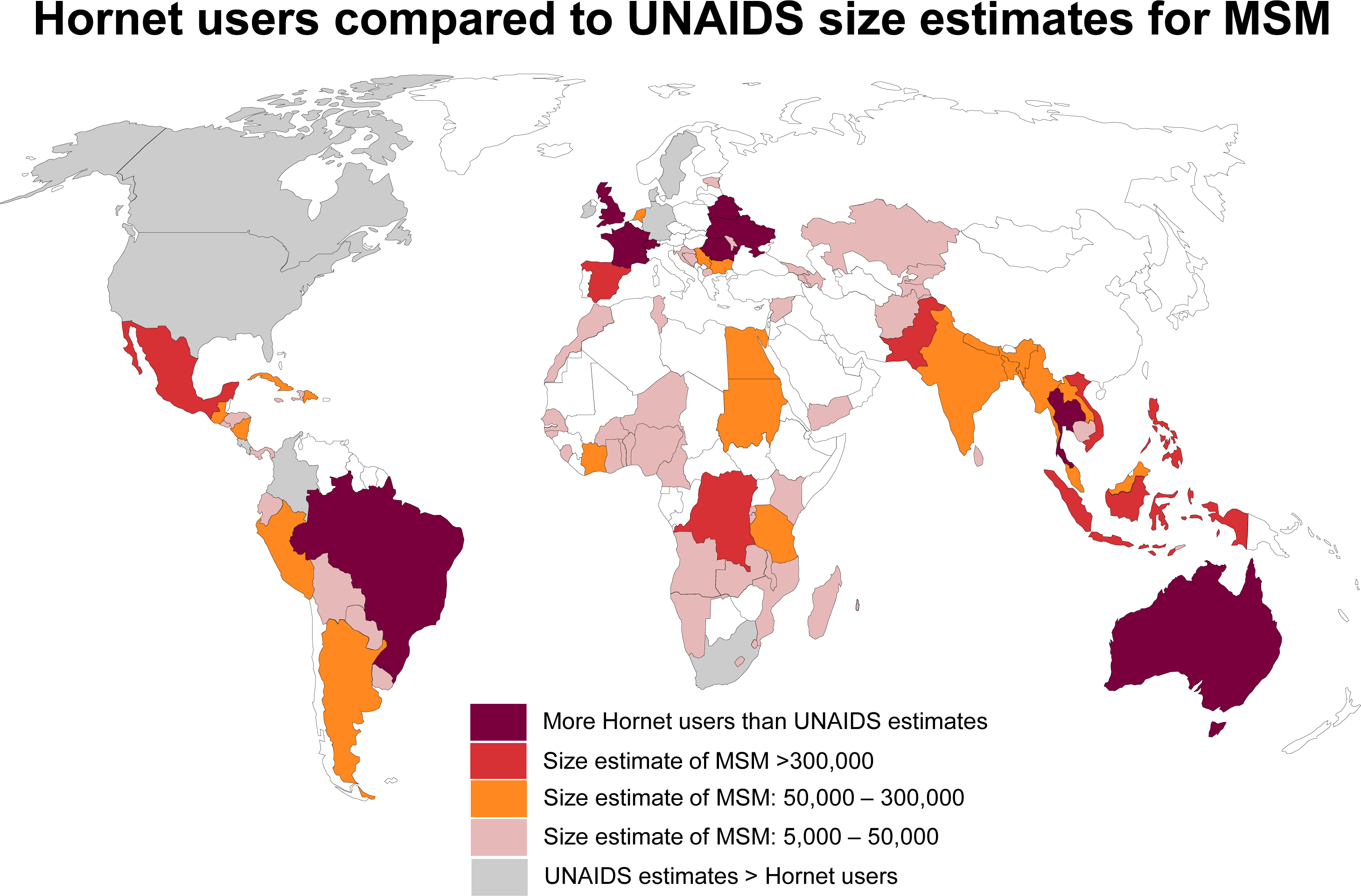
In France, Thailand and Romania, Hornet users are double than the UNAIDS estimates. White represents countries with no data orr <5,000 MSM. The size estimates of MSM (for countries in red, orange and pink) are from UNAIDS AIDS info Online Database and PEPFAR COP

## Results

Comparing UNAIDS with dating application user data revealed a number of discrepancies (Table 1). Of the 29 comparison countries, only 15 had UNAIDS or other MSM population estimates in the public domain. In 9 of 29 countries, the number of gay dating application users were above the UNAIDS estimate of the number of MSMs with 8 over 30% higher and three more than double the reported estimate. In 9 countries UNAIDS estimates were larger than dating application user numbers, perhaps reflecting early or incomplete market penetration for the application. Gay dating application users are younger and tech savvy and likely represent a sub-set of all MSM‐‐the application user totals are probably an underestimate of the total MSM population. For the 29 countries, we found only 7 partial or complete nationally representative care continua for MSM published between 2010 and 2016. Among estimated MSM living with HIV, viral suppression varied between 42% (United States) to 99% (Denmark). The quality of the continua methods varied (quality data not shown) (18-20).

**Table 1:**
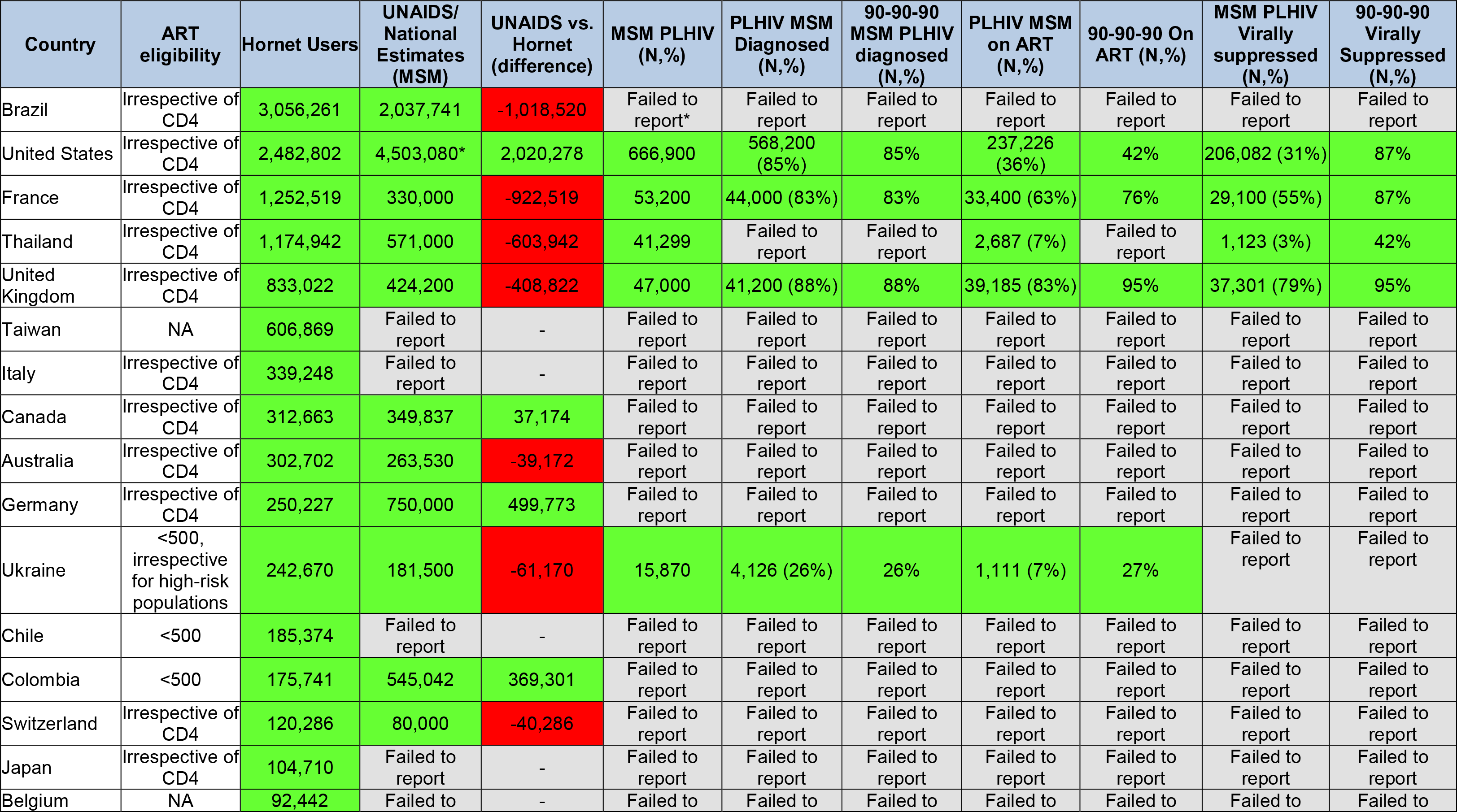
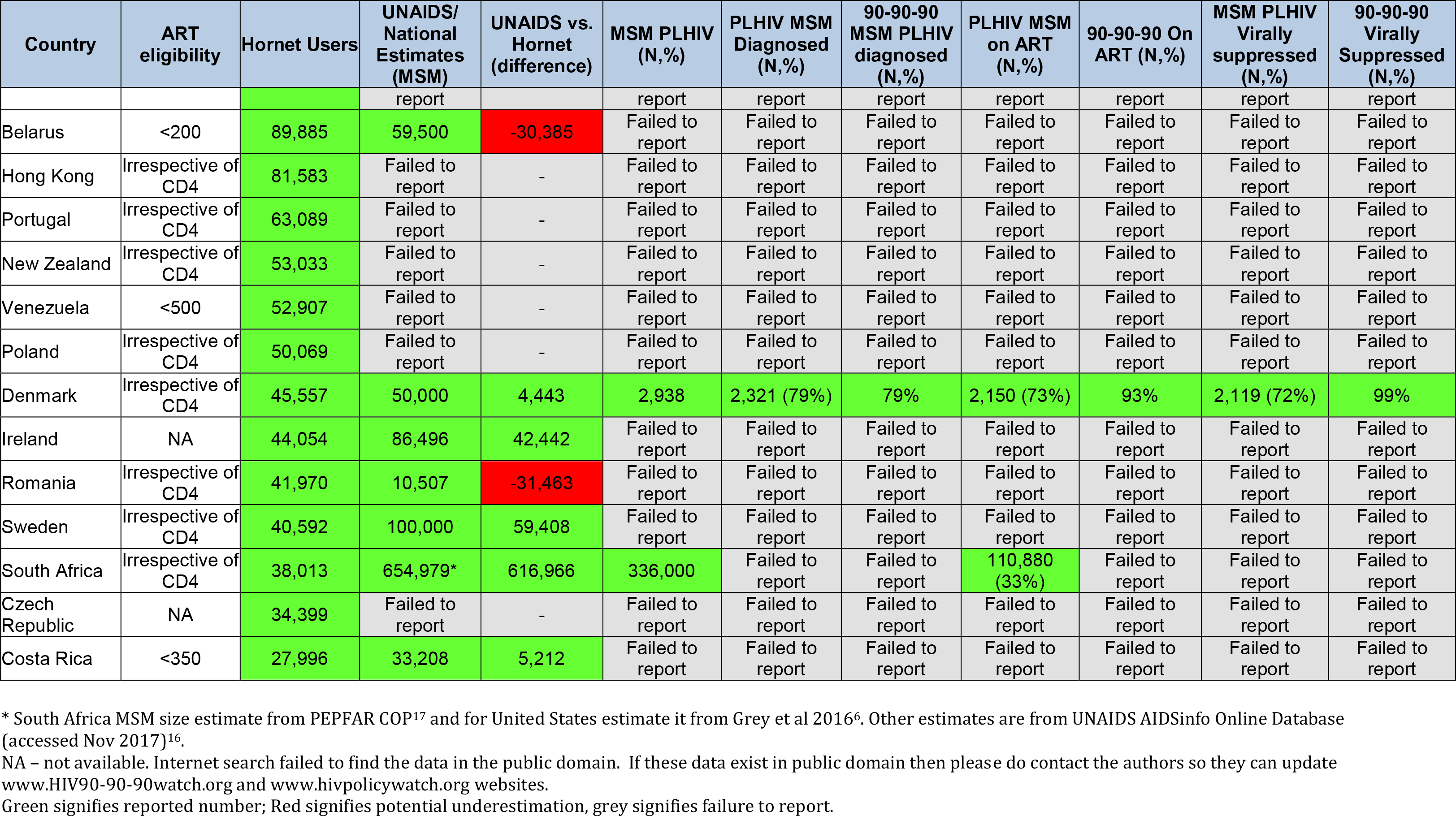
MSM:Hornetusers,MSMestimates,90-90-90andcarecontinua.

## Discussion

The response to HIV has included unprecedented advances in our knowledge and understanding of how to end the epidemic. Access to HIV services including care, treatment and other prevention interventions has saved millions of lives. Since 2003, global efforts to scale up antiretroviral therapy (ART) have played a major role in halting and reversing the HIV epidemic in many settings. Recent scientific advances have demonstrated that immediate use of ART irrespective of CD4 cell count can reduce HIV-related morbidity, mortality, transmission and costs (21-24). However, for the benefits of immediate ART to be fully realized, people living with HIV (PLHIV), especially the high-risk populations, must optimally engage to support each step along the HIV care continua (25), from early diagnosis to access to ART and viral suppression. Our study highlights significant discrepancies between official estimates of MSM and the number of individual users of gay dating applications. While only a selective sample of gay dating application users, 30% higher numbers in 8 countries suggest that the current UNAIDS and national methodology for estimating the number of MSM should be reconsidered in light of this new data source.

Our study has potential limitations including uncertainties and variations inherent in the UNAIDS MSM estimation process, the paucity of published national cascades for key populations, and the potential to only temporarily use the gay dating application to meet other men. Additionally, given the existence of multiple gay dating applications, relying on data from only one application could result in an underestimation of MSM who may be using another application. Finding higher gay dating application user numbers when compared with official estimates makes common sense since it is arguably easier and more anonymous to use a gay dating application than to respond to interview questions from clinicians or people implementing household or other official or research surveys. While certainly not definitive, the number of people using gay dating applications provides another potential data point that can be used when considering the HIV response strategy including service delivery efforts for MSM.

“What is not monitored is not done” and global estimates of key population denominators are vital to a successful HIV response. UNAIDS leads the global estimates working group that develops national estimates for key populations (16). While some have made the legitimate argument that collecting data on MSM can put their lives at risk in some settings, it is also recognized that without collecting this information it will be difficult to know whether MSM are successfully accessing services. Additionally, this lack of information hampers the ability to design and properly fund MSM specific services. Our study showed significant differences between the number of users of a popular gay dating application when compared with UNAIDS estimates for some countries. There are many possible reasons for this including lack of national studies or data, inaccurate modeling, unknown HIV prevalence, incidence and/or treatment access for MSM. Additionally, some countries deny the existence of MSM. Regardless, the social media derived data from Gay dating applications provide an important insight into population estimates and could be used in estimation exercises. Estimates are just that, estimates, and information that provides a means to triangulate on data points should help improve their accuracy. Consideration could also be given to collecting information about service access via social media as a means to triangulate on service delivery and quality. The global estimates group reliance on traditional surveillance techniques appears to result in inaccurate denominators and the inability to accurately measure progress towards HIV services access for vulnerable populations including MSM.

The care continua have become vital tools to monitor the HIV programmatic efforts and guide future actions to achieve the 90-90-90 target by 2020 or 73% of the estimated PLHIV virally suppressed along the continua (1, 2). Discrimination, stigma, and lack of data regarding denominators are all obstacles to successful delivery of high quality services. HIV care programs specifically tailored for MSM, which include community-based HIV testing and immediate access to ART and viral load monitoring, need to be scaled-up in partnership with civil society organizations. This will not be possible without addressing the social and legal barriers impeding access to care for key populations. Additionally, more and better quality data on key populations is needed in the public domain to promote accountability and data-driven programs. Countries and organizations working with key populations need to strengthen their current monitoring and evaluation (M&E) systems so that more real-time, local and disaggregated programmatic data can be collected and continua can be standardized across countries. Investment in population-based surveys, targeted surveys in areas with high HIV prevalence among key populations, case-based reporting and unique identification codes can help countries construct more reliable continua for key populations (26). Community engagement will be critical in both the design and implementation of M&E activities to ensure a human rights framework is followed and that both communities and health authorities can own the data and work together to improve access to diagnosis and treatment. Social media, with its extraordinary reach, has considerable potential to improve the quality of the HIV response.

Achieving 90-90-90 is vital for MSM to prevent illness, death and transmission. However, 90-90-90 translates into 27% of PLHIV not on ART and virally suppressed. Knowing and being able to estimate the number of people who may be at risk is a fundamental element of public health and disease control. Gay dating applications and other social media platforms provide unique insights into how to reach and include people in solving the HIV problem. Gay dating applications can provide a powerful platform for peer-to-peer sharing of health information and delivering HIV-related services (27, 28) while also helping public health efforts by providing insights regarding the size and characteristics of the community. Social media has significant promise and may help provide the best information available to ensure that MSM and other vulnerable people living with HIV and their communities are not left behind on the way to ending AIDS.

## Footnotes

1 Here we use MSM but the community prefers the word gay

